# Scrutinization of Soil Seed Bank from Arid to Mesic Habitats of Dera Ghazi Khan

**DOI:** 10.1101/333856

**Authors:** Allah Bakhsh Gulshan, Ali Bakhsh, Syed Mazhar Irfan, Sabir Hussain, Khazir Hayat, Muhammad Imran Atta

**Author notes:** Corresponding author: Name: Allah Bakhsh Gulshan, Associate Professor of Botany Department of Botany/ Director Quality Enhancement, Cell Ghazi University Dera Ghazi Khan Punjab Pakistan, Email ID or.

## Abstract

The research was conducted to investigate the persistent soil seed bank composition and its relation to the above-ground vegetation of upland area (piedmont) to low land area (alluvial) landscape from arid to mesic region of Dera Ghazi Khan. A transact of 40 kilometers was laid down from arid to mesic habitat. At each 2 km a quadrate of 1 m^−2^ sizes was thrown in the field to collect a soil sample of 2kg from soil cores ranging 0-15 cm deep for the analysis of soil seed bank. Twenty different sites were sampled by throwing 6 quadrates at each site making a total of 120 samples. Three thousand seeds were obtained of 50 different species from all the collected samples. Soil seed bank density m^−2^ was higher in the alluvial plains of Dera Ghazi Khan. Most of the perennial species, which were xerophytic in nature such as *Aerua persica, Calotropis procera, Fagonia indica, Leptadaenia pyrotechnica, Peganum hermala, Rhazya stricta* and *Suaeda fructicosa* were found in the piedmont (arid) soil habitat and the soil seed bank relatively less than the species found in the alluvial (Mesic) soil habitat, which were mostly of annual life span such as *Chenopodium murale, Euphorbia prostrata, Medicago denticulata, Fumaria indica*, and *Withania somnifera*. From this study it is concluded that the similarity found between soil seed bank and above ground vegetation of both historic types of habitats piedmont (arid) and alluvial (mesic) of Dera Ghazi Khan

## Introduction

The subtropical region of Pakistan in general and the special area of Dera Ghazi Khan consist of diversified land habitats. Due to this inclined demarcation in the land from west-east the above ground vegetation is certainly the shadow of below ground vegetation. The significance of soil seed banks and their role in the natural diversity of plant species of many ecosystems is well-recognized various research across the globe like Young et al., (1987); Garwood, (1989); Forcella et al., (1992); Adams et al., (2005); Tang et al., (2006). Moreover, Miao, et al., (2016) similarity index between soil seed bank and ground vegetation was explored in sandy environment. Soil seed banks function as natural seed reserves for future regeneration of many plant species, yet have only recently been incorporated into demographic models for plant population monitoring. The reasons for this are in large part due to the difficulty in collecting seed bank data. Arable and non arable area of soil in this region about soil seed banks were first studied intensively because of their important economic impacts in agricultural systems Davis et al., (2005); Chee-Sanford et al., (2006). The seeds present in the profile of soil play a very important role in the prediction of above ground vegetation and similar type of studies were performed by Luzuriaga et al. (2005). But a large group of researchers like Chippindale and Milton, (1934); Champness and Morris, (1948); Major and Pyott, (1966); Thompson and Grime, (1979) were believed that the below ground vegetation were totally different from the above ground vegetation. The gap of this controversial analysis about the soil seed bank is a real attribute, because the size of above ground vegetation in somehow the reflection of below ground vegetation or not, this is totally depending on the regional climatic and edaphic factors of the area. The actual picture about the soil seed bank and their relationship to above round vegetation is still remaining debatable for a long time, because such type of variations varies with the variation of spatial and temporal changes link to natural factors. Similarity and dissimilarity view point about the about ground vegetation and below ground vegetation in terms of soil seed bank were available in the literature across the globe e.g. Thompson and Grime, (1979); Grime et al., (1988); Pierce and Cowling, (1991); Mountford et al., (1993a, b); Milberg (1995); Bakker et al., (1996); Kirkham et al., (1996); Kirkham and Kent, (1997). Chang et al., (2001); Caballero et al., (2003); Caballero et al., (2008) Thompson and; Hodgson et al., (1995), all were the idea and observed that the above ground vegetation closely resembles with the below ground vegetation in terms of soil seed bank. In other side Fenner and Thompson (2005) and Esmeralda et al., (2011), they were followed the idea of dissimilarity concept about soil seed bank and above round vegetation. Analysis of soil seed bank characterization in different substrates were previously investigated through scholars like He et al., (2015, 2016) by applying advanced ecological methods.

The distribution of vegetation in reference to space and soil seed bank was found significant relationship among one another and this evidence supported the work of Navarra and Quintana-Ascencio, (2012). Others workers group like Milberg (1995); Bakker et al. (1996); Peco et al. (1998); Ferrandis et al. (2001); Figueroa et al. (2004); Caballero et al. (2005); Caballero et al., (2008) and Martinez-Duro et al., (2010) were supported the document of similarity found among the soil seed bank and above ground vegetation. But Kemp (1989) was not supported the similarity relationship b/w seed bank and vegetation. According the observation of Morin and Payette the similarity was found in same place/habitat but not to another habitat Moreover analogous type of correlation were observed in different types of habitats by the large number of workers such as Cabin et al., (2000), in alpine habitats; Whipple, (1978); Diemer and Prock, (1993); Ingersoll and Wilson, (1993); Stanifforth et al., (1998), in Wetlands environments Leck and Graveline, (1979); Parker and Leck, (1985); Leck and Simpson, (1995) and in Salt pans Ungar and Riehl, (1980). The invasive species were continually changed the community of the area and also to become the part of soil seed bank as an invaded species according to Gioria and Pysek (2016).

The main objectives of this study to explore and predict the soil seed bank and their relationships with above ground vegetation along the inclined gradient landscape from upland area (piedmont arid environment) to low land (alluvial mesic environment) plains of Dera Ghazi Khan.

## Materials & Methods

Sampling data of seed bank was recorded concomitantly from the selected sites for study area. Seed bank samples were collected in March-April and September-October in the year 2017. For seed bank five 1 m^−2^ quadrates were used in each sites and a total of 60 sites. Samples were containing five soil cores. Each soil core has 4 cm in diameter with 15 cm deep in length from surface in vertical direction. The total area sampled at each site was 0.125m^2^. Sampling sites were shown in Figure 1.

**Figure 1.**
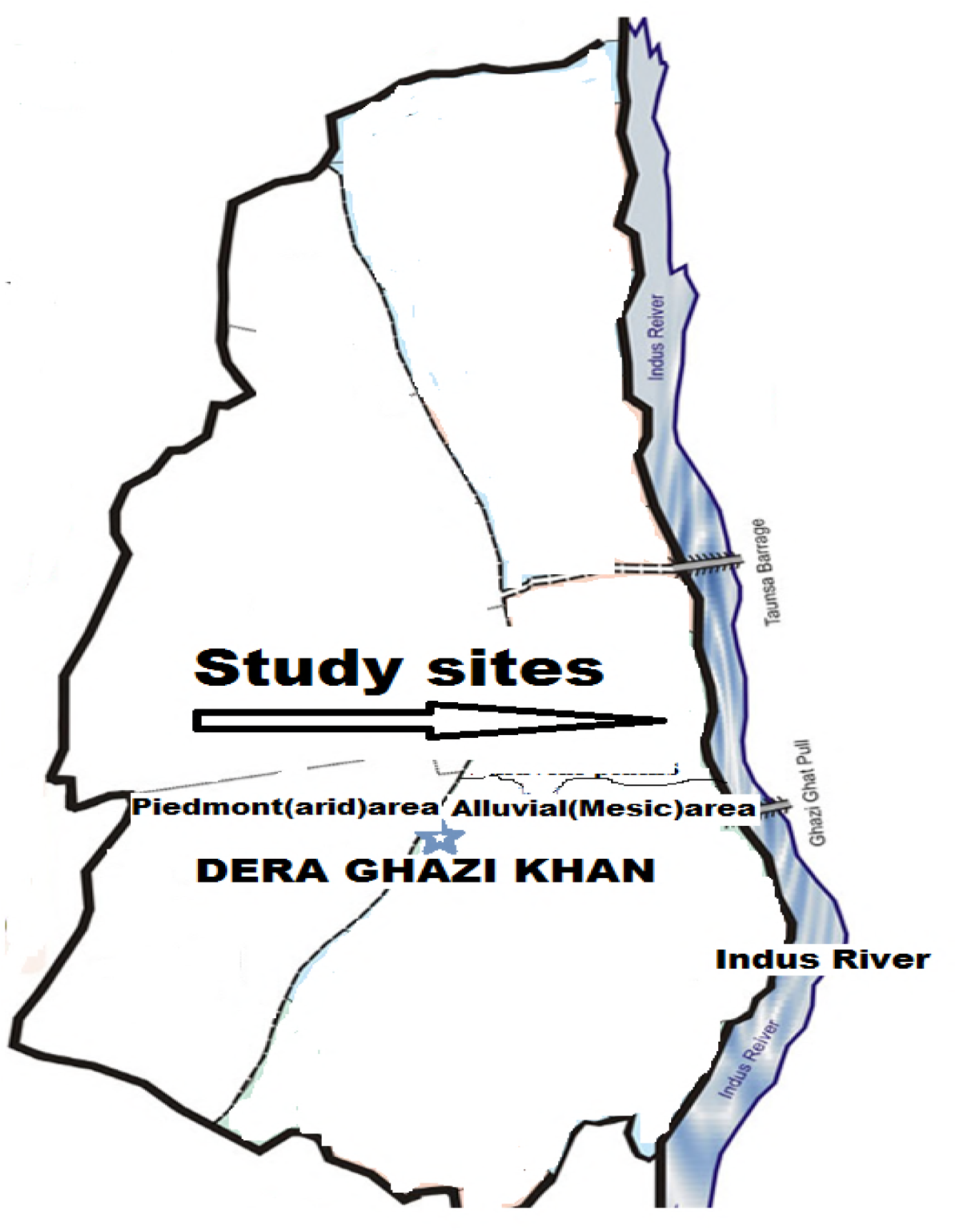
Study area mentioned in selected box and arrow indicates the sampling units started from piedmont (aria) o alluvial (mesic) plains.

The collected samples for soil seed bank were treated by followed the methods of Ter Heerdt et al., (1996) and Bekker et al., (2000). A fine sieve ranges size (mesh width 0.212mm) were used to wash the seeds from soil samples before being spread out in a very thin layer (< 3mm) on trays filled with sterile potting soil mix with fine sand and allowed to germinate in a glasshouse for 8 weeks’ duration. The trays were kept under a 12 hours’ photoperiod with night and day temperatures 24/30°C and watering after 24 hours daily. Wait for the total emergence of seeds and counted germination on daily basis, when there was no further emergence and all seedlings had been identified, counted and removed, the samples were recollected from the trays, dried and stored at room temperature and dark until sorted for the remaining seeds. The germination method was the best one for exploring and quantifying the viable seeds form the collected soil samples and seedling emergence methods to assess the soil seed bank was followed by Falinska, (1998,1999), Weiterova, (2008), Koch et al., (2011). The lesser no of seeds which have to fail in germination and these were excluded from the present analysis.

### Collection of seed bank data

The soil seed bank data were sampled from four habitats in *Valley piedmont rain-fed sites, Piedmont plains sites*, western bank of Indus River and sites along eastern bank of Indus River. Sampling area were chosen a priori based on the moist gradient i.e. proximity from the river bank. *Rain-fed* valley sites are much away from the river with xeric environment while the alluvial sites are much closer with mesophytic habitat. The detailed description of the habitats under investigation is given in table (1).

**Table 1.** Description and details of sampling sites from which soil seed bank data was recorded in year 2017. Each sampling sites consists of > 0.1-hectare area.

| Study sites | No. of sites | Area survey (km) | Proximity from river (km) | Description of sites |
| --- | --- | --- | --- | --- |
| Valley piedmont<br>Rain-fed sites | 15 | 6 | 30-40 | Here no regular cultivation. the area is irrigated almost through hill torrents |
| Piedmont plains areas | 15 | 6 | 20-25 | The area is irrigated through canal water and almost the soil is arable |
| Western alluvial sites | 15 | 6 | 15-20 | The area is consist of arable land and cultivated with seasonal crops regularly |
| Eastern alluvial sites | 15 | 6 | 5-10 | The area is consist of arable land and cultivated with seasonal crops regularly and irrigated through canal water cum tubewell (underground water) |

## RESULTS

In the current scenario of study was the analysis of three microhabitats of soil for the seed bank. A total of 50 species in soil seed bank were scrutinized from the four microhabitats of piedmont (arid) to alluvial (mesic) inclined area of Dera Ghazi Khan. The recovered soil seed bank of different plant species from two major (piedmont and alluvial) soil habitats indicated that lesser number of soil seed bank (890±3.53) was recovered from the piedmont site which was two folds lesser than the alluvial sites. The highest number of seed bank (2110±14.21) was scrutinized from the soil of alluvial plains of Dera Ghazi Khan (Table 2).

**Table 2.** Total recovered seeds of various plant species from four different soil habitats of Dera Ghazi Khan.

| Study sites | No. of sites | Area survey (km) | Proximity from river (km) | Total Soil Seed Bank recovered from each habitat |  |
| --- | --- | --- | --- | --- | --- |
|  |  |  |  | Number of species | Total recovered soil seed bank |
| Valley piedmont Rain-fed sites | 15 | 6 | 30-40 | 08 | 230 |
| Piedmont plains areas | 15 | 6 | 20-25 | 14 | 660 |
| Western alluvial sites | 15 | 6 | 15-20 | 22 | 560 |
| Eastern alluvial sites | 15 | 6 | 5-10 | 30 | 1550 |
| **4 major soil habitats** | **60** | **24** | **40** | **74** | **3000 (Total recovered seeds)** |

\* Total 50 different species soil seed banks recovered from 4 major soil habitats of Dera Ghazi Khan, although 24 different common species of plants and their soil seeds recovered almost from all the 4 habitats

The perennial life span species such as, *Fagonia indica, Peganum hermala, Rhazya stricta, Suaeda fructicosa, Tribulus terristris* and *Withania coagulens* were the abundant species in relation to soil seed bank of the upland (arid) area of Dera Ghazi Khan. But the richness of annual life span species such as *Chenopodium album, Fumaria indica, Medicago denticulata* and *Melilotus albus* were found maximum in the soil seed bank of alluvial flat plains (mesic) area (Table 3). The soil seed bank belonging to xeric type of vegetation was abundant in piedmont (upland area) whereas the mesic type of soil seed bank was mostly recovered from the alluvial plain (low land area). Table 3 clearly indicates the occurrence of various species in terms of rare, frequent, abundant and vary abundant in relation to soil seed bank. Above ground vegetation was the mirror image of below ground vegetation. *Acacia modesta* was frequently found in piedmont area and *Acacia nilotica* was found in alluvial plains of Dera Ghazi Khan. *Euphorbia helioscopia, Fumaria indica, Melilotus albus* and *Vicia sativa* are the very abundant species of mesic habitats.

**Table 3.** Species occurring locality wise in vegetation (Veg.) or in the soil seed bank (S.B.), with (V.A. = very abundance, A. = abundance, F. = frequent and R. = rare), two different Localities of Dera Ghazi Khan.

| S. No. | Species Name | Piedmont soil environment |  | Alluvial plains soil environment |  |
| --- | --- | --- | --- | --- | --- |
|  |  | Soil seed bank | Vegetation | Soil seed bank | Vegetation |
| 1 | *Abutilon* | 12.00 | R | 40.00 | F |
|  | *indicum* |  |  |  |  |
| 2 | *Acacia modesta* | 28.00 | F | - | - |
| 3 | *Acacia nilotica* | 40.00 | F | 8.00 | R |
| 4 | *Achyranthes aspera* | 8.0 | R | 18 | F |
| 5 | *Aerua persica* | 60.00 | A | - | - |
| 6 | *Alhagi maurorum* | - | - | 80 | A |
| 7 | *Amaranthus viridis* | - | - | 40 | F |
| 8 | *Brachiaria ramosa* | 14.00 | R | 78.00 | A |
| 9 | *Calotropis procera* | 35.00 | F | 6.00 | R |
| 10 | *Capparis aphylla* | 18.00 | F | 6.00 | R |
| 11 | *Chenopodium album* | - | - | 204.00 | V.A |
| 12 | *Chenopodium murale* | 8.00 | R | 86.00 | A |
| 13 | *Chrozophora tinctoria* | - | - | 32.00 | F |
| 14 | *Citrus colocynthis* | 30.00 | F | - | - |
| 15 | *Conyza bonariensis* | - | - | 6.00 | R |
| 16 | *Cronopus didynamus* | - | - | 8.00 | R |
| 17 | *Cyperus rotundus* | - | - | 8.00 | R |
| 18 | *Dalbergia sisso* | - | - | 10.00 | F |
| 19 | *Datura metel* | 11 | R | - | - |
| 20 | *Datura stramonium* | - | - | 16 | R |
| 21 | *Euphorbia helioscopia* | - | - | 165.00 | V.A |
| 22 | *Euphorbia prostrata* | 6.00 | R | 44.00 | A |
| 23 | *Fagonia indica* | 74.00 | A | 8.00 | R |
| 24 | *Fumaria* | - | - | 229.00 | V.A |
|  | *indica* |  |  |  |  |
| 25 | *Heliotropium strigosum* | 18.00 | R | 24.00 | F |
| 26 | *Lathyrus aphaca* | - | - | 126.00 | V.A |
| 27 | *Leptadaenia pyrotechnica* | 22.00 | F | - | - |
| 28 | *Medicago denticulata* | - | - | 108.00 | V.A |
| 29 | *Melilotus albus* | - | - | 266.00 | V.A |
| 30 | *Oxalis corniculata* | - | - | 4.00 | R |
| 31 | *Panicum antidotale* | 8.00 | R | - | - |
| 32 | *Peganum harmala* | 74.00 | A | - | - |
| 33 | *Phalaris minor* | - | - | 64.00 | F |
| 34 | *Portulaca oleracea* | - | - | 40.00 | F |
| 35 | *Pulicaria crispa* | - | - | 42 | F |
| 36 | *Ranunculus muricatus* | - | - | 34 | F |
| 37 | *Ranunculus scleratus* | - | - | 26 | F |
| 38 | *Rhazya stricta* | 86.00 | V.A | - | - |
| 39 | *Rumex dentatus* | - | - | 58.00 | F |
| 40 | *Sonchus asper* | - | - | 3.00 | R |
| 41 | *Solanum nigrum* | - | - | 40.00 | A |
| 42 | *Solanum surrattense* | 65.00 | A | - | - |
| 43 | *Sorghum halepense* | - | - | 36.00 | F |
| 44 | *Sophora mollis* | - | - | 34.00 | F |
| 45 | *Suaeda fruticosa* | 80.00 | A | - | - |
| 46 | *Tribulus terrestris* | 48.00 | F | 24.00 | R |
| 47 | *Trigonella corniculata* | - | - | 65.00 | A |
| 48 | *Vicia sativa* | - | - | 103.00 | V.A |
| 49 | *Withania coagulens* | 78.00 | A | - | - |
| 50 | *Withania somnifera* | - | - | 60.00 | F |

### Soil seed bank analyzed from different depths

Soil preserves the seed bank in its micelles at various depths. Maximum soil seed bank was explored from the depth of surface to ten centimeter depths. When the depth of soil increased then the recovery of soil seed bank from the soil micelles was lesser. The fresh and almost annual life span species seed bank was put down at the depth of surface to 5cm (Table 4). Light weight such as *Chenopodium sp, Fumaria indica and Melilotus* sp etc and minute with assisted seeds including *Calotropis procera, Conyza bonariensis* etc were mostly present at this depth. The contribution of larger seeds species in the soil seed bank such as *Acacia sp, Alhagi maurorum* and *Dulbergia sisso* etc were available more or less at the various depths from surface to 15cm of soil profiles. Maximum number of soil seed bank was recovered from the alluvial soil or the mesic plain areas of the Dera Ghazi Khan, where all types of vegetation are cultivated annually (Table 4).

**Table 4.**
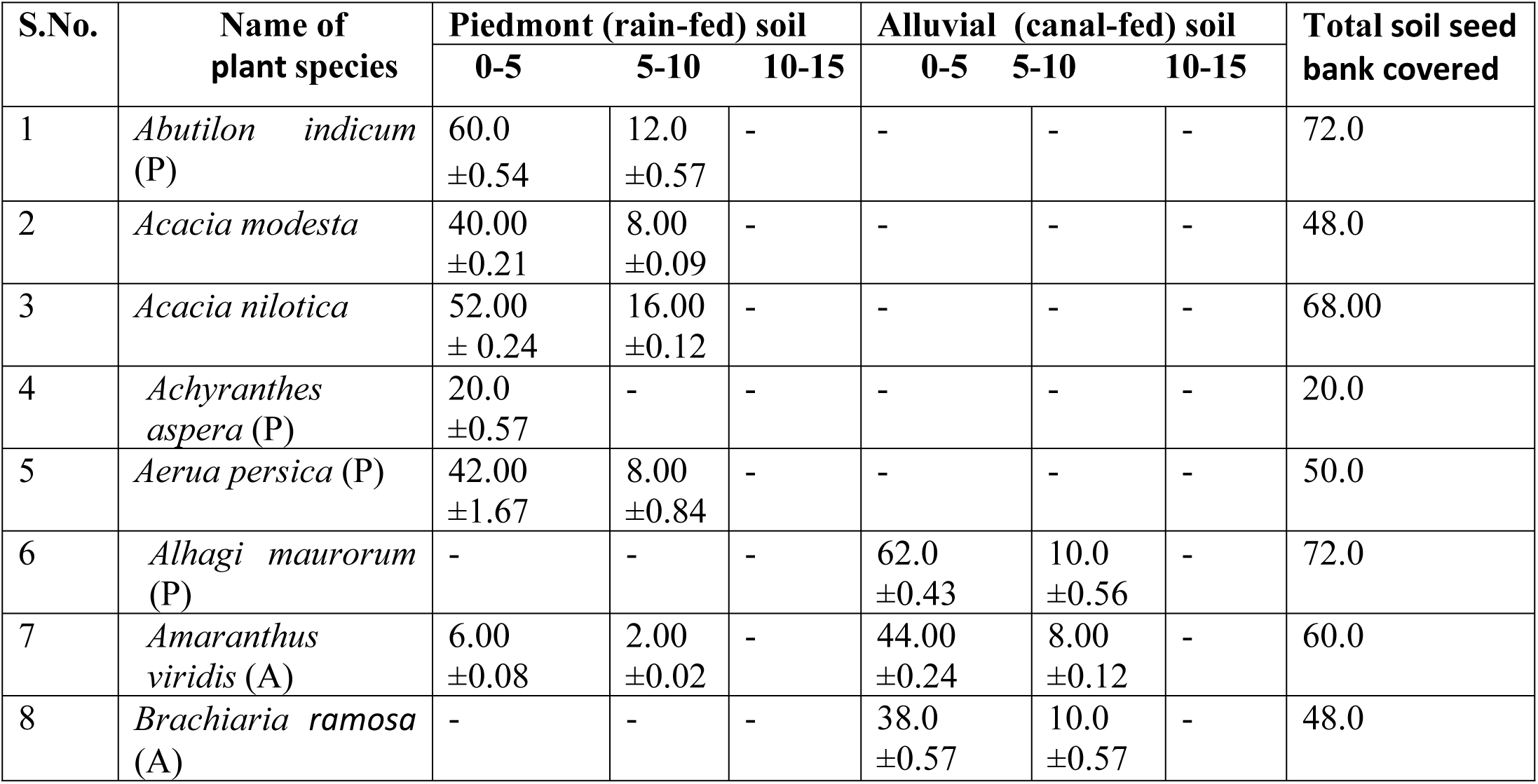
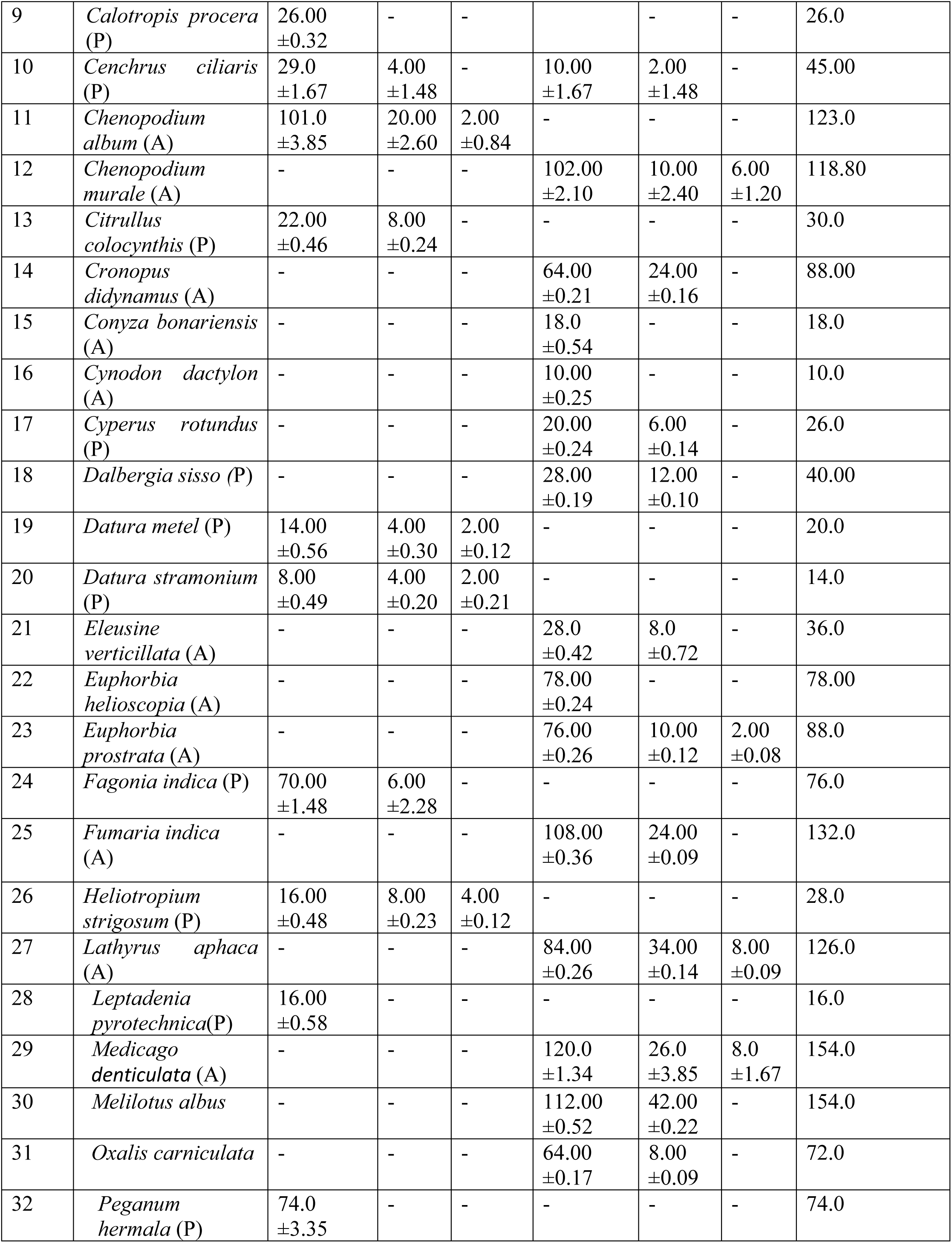
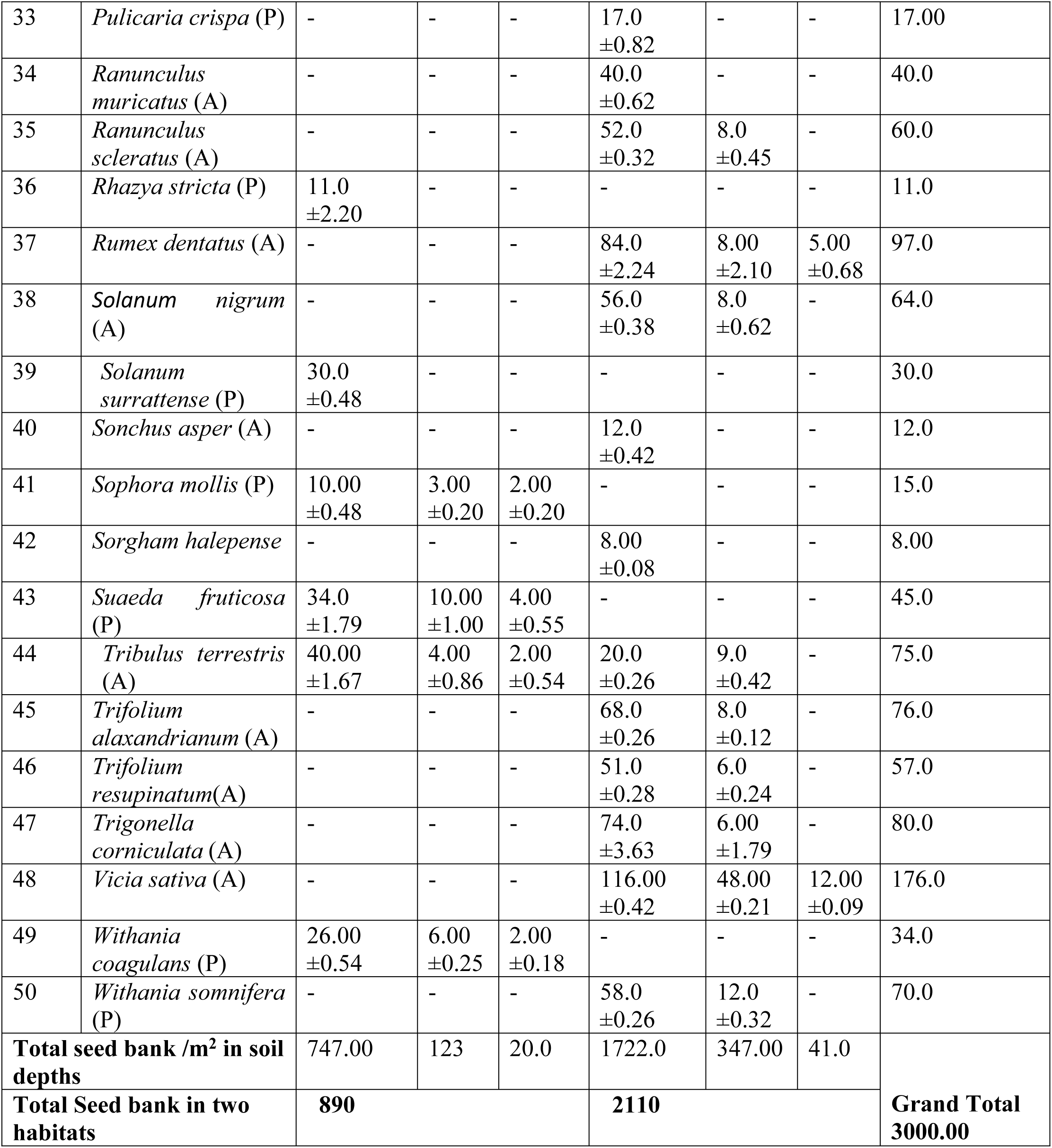
Soil seed bank densities of species (m^−2^) in two different habitats (Piedmont & Alluvial soil). Each sample contains 5 pooled soil cores. Samples were taken in to three different soil strata (0-5cm; 5-10 cm; 10-15 cm). Annual (A) Perennial (P)

| S.No. | Name of plant species | Piedmont (rain-fed) soil |  |  | Alluvial (canal-fed) soil |  |  | Total soil seed bank covered |
| --- | --- | --- | --- | --- | --- | --- | --- | --- |
|  |  | 0-5 | 5-10 | 10-15 | 0-5 | 5-10 | 10-15 |  |
| 1 | *Abutilon indicum* (P) | 60.0<br>±0.54 | 12.0<br>±0.57 | - | - | - | - | 72.0 |
| 2 | *Acacia modesta* | 40.00<br>±0.21 | 8.00<br>±0.09 | - | - | - | - | 48.0 |
| 3 | *Acacia nilotica* | 52.00<br>± 0.24 | 16.00<br>±0.12 | - | - | - | - | 68.00 |
| 4 | *Achyranthes aspera* (P) | 20.0<br>±0.57 | - | - | - | - | - | 20.0 |
| 5 | *Aerua persica* (P) | 42.00<br>±1.67 | 8.00<br>±0.84 | - | - | - | - | 50.0 |
| 6 | *Alhagi maurorum* (P) | - | - | - | 62.0<br>±0.43 | 10.0<br>±0.56 | - | 72.0 |
| 7 | *Amaranthus viridis* (A) | 6.00<br>±0.08 | 2.00<br>±0.02 | - | 44.00<br>±0.24 | 8.00<br>±0.12 | - | 60.0 |
| 8 | *Brachiaria ramosa* (A) | - | - | - | 38.0<br>±0.57 | 10.0<br>±0.57 | - | 48.0 |
| 9 | *Calotropis procera* (P) | 26.00<br>±0.32 | - | - |  | - | - | 26.0 |
| 10 | *Cenchrus ciliaris* (P) | 29.0<br>±1.67 | 4.00<br>±1.48 | - | 10.00<br>±1.67 | 2.00<br>±1.48 | - | 45.00 |
| 11 | *Chenopodium album* (A) | 101.0<br>±3.85 | 20.00<br>±2.60 | 2.00<br>±0.84 | - | - | - | 123.0 |
| 12 | *Chenopodium murale* (A) | - | - | - | 102.00<br>±2.10 | 10.00<br>±2.40 | 6.00<br>±1.20 | 118.80 |
| 13 | *Citrullus colocynthis* (P) | 22.00<br>±0.46 | 8.00<br>±0.24 | - | - | - | - | 30.0 |
| 14 | *Cronopus didynamus* (A) | - | - | - | 64.00<br>±0.21 | 24.00<br>±0.16 | - | 88.00 |
| 15 | *Conyza bonariensis* (A) | - | - | - | 18.0<br>±0.54 | - | - | 18.0 |
| 16 | *Cynodon dactylon* (A) | - | - | - | 10.00<br>±0.25 | - | - | 10.0 |
| 17 | *Cyperus rotundus* (P) | - | - | - | 20.00<br>±0.24 | 6.00<br>±0.14 | - | 26.0 |
| 18 | *Dalbergia sisso* (P) | - | - | - | 28.00<br>±0.19 | 12.00<br>±0.10 | - | 40.00 |
| 19 | *Datura metel* (P) | 14.00<br>±0.56 | 4.00<br>±0.30 | 2.00<br>±0.12 | - | - | - | 20.0 |
| 20 | *Datura stramonium* (P) | 8.00<br>±0.49 | 4.00<br>±0.20 | 2.00<br>±0.21 | - | - | - | 14.0 |
| 21 | *Eleusine verticillata* (A) | - | - | - | 28.0<br>±0.42 | 8.0<br>±0.72 | - | 36.0 |
| 22 | *Euphorbia helioscopia* (A) | - | - | - | 78.00<br>±0.24 | - | - | 78.00 |
| 23 | *Euphorbia prostrata* (A) | - | - | - | 76.00<br>±0.26 | 10.00<br>±0.12 | 2.00<br>±0.08 | 88.0 |
| 24 | *Fagonia indica* (P) | 70.00<br>±1.48 | 6.00<br>±2.28 | - | - | - | - | 76.0 |
| 25 | *Fumaria indica* (A) | - | - | - | 108.00<br>±0.36 | 24.00<br>±0.09 | - | 132.0 |
| 26 | *Heliotropium strigosum* (P) | 16.00<br>±0.48 | 8.00<br>±0.23 | 4.00<br>±0.12 | - | - | - | 28.0 |
| 27 | *Lathyrus aphaca* (A) | - | - | - | 84.00<br>±0.26 | 34.00<br>±0.14 | 8.00<br>±0.09 | 126.0 |
| 28 | *Leptadenia pyrotechnica* (P) | 16.00<br>±0.58 | - | - | - | - | - | 16.0 |
| 29 | *Medicago denticulata* (A) | - | - | - | 120.0<br>±1.34 | 26.0<br>±3.85 | 8.0<br>±1.67 | 154.0 |
| 30 | *Melilotus albus* | - | - | - | 112.00<br>±0.52 | 42.00<br>±0.22 | - | 154.0 |
| 31 | *Oxalis corniculata* | - | - | - | 64.00<br>±0.17 | 8.00<br>±0.09 | - | 72.0 |
| 32 | *Peganum harmala* (P) | 74.0<br>±3.35 | - | - | - | - | - | 74.0 |
| 33 | *Pulicaria crispa* (P) | - | - | - | 17.0<br>±0.82 | - | - | 17.00 |
| 34 | *Ranunculus muricatus* (A) | - | - | - | 40.0<br>±0.62 | - | - | 40.0 |
| 35 | *Ranunculus scleratus* (A) | - | - | - | 52.0<br>±0.32 | 8.0<br>±0.45 | - | 60.0 |
| 36 | *Rhazya stricta* (P) | 11.0<br>±2.20 | - | - | - | - | - | 11.0 |
| 37 | *Rumex dentatus* (A) | - | - | - | 84.0<br>±2.24 | 8.00<br>±2.10 | 5.00<br>±0.68 | 97.0 |
| 38 | *Solanum nigrum* (A) | - | - | - | 56.0<br>±0.38 | 8.0<br>±0.62 | - | 64.0 |
| 39 | *Solanum surrattense* (P) | 30.0<br>±0.48 | - | - | - | - | - | 30.0 |
| 40 | *Sonchus asper* (A) | - | - | - | 12.0<br>±0.42 | - | - | 12.0 |
| 41 | *Sophora mollis* (P) | 10.00<br>±0.48 | 3.00<br>±0.20 | 2.00<br>±0.20 | - | - | - | 15.0 |
| 42 | *Sorgham halepense* | - | - | - | 8.00<br>±0.08 | - | - | 8.00 |
| 43 | *Suaeda fruticosa* (P) | 34.0<br>±1.79 | 10.00<br>±1.00 | 4.00<br>±0.55 | - | - | - | 45.0 |
| 44 | *Tribulus terrestris* (A) | 40.00<br>±1.67 | 4.00<br>±0.86 | 2.00<br>±0.54 | 20.0<br>±0.26 | 9.0<br>±0.42 | - | 75.0 |
| 45 | *Trifolium alaxandrianum* (A) | - | - | - | 68.0<br>±0.26 | 8.0<br>±0.12 | - | 76.0 |
| 46 | *Trifolium resupinatum*(A) | - | - | - | 51.0<br>±0.28 | 6.0<br>±0.24 | - | 57.0 |
| 47 | *Trigonella corniculata* (A) | - | - | - | 74.0<br>±3.63 | 6.00<br>±1.79 | - | 80.0 |
| 48 | *Vicia sativa* (A) | - | - | - | 116.00<br>±0.42 | 48.00<br>±0.21 | 12.00<br>±0.09 | 176.0 |
| 49 | *Withania coagulans* (P) | 26.00<br>±0.54 | 6.00<br>±0.25 | 2.00<br>±0.18 | - | - | - | 34.0 |
| 50 | *Withania somnifera* (P) | - | - | - | 58.0<br>±0.26 | 12.0<br>±0.32 | - | 70.0 |
| Total seed bank /m² in soil depths |  | 747.00 | 123 | 20.0 | 1722.0 | 347.00 | 41.0 | Grand Total<br>3000.00 |
| Total Seed bank in two habitats |  | 890 |  |  | 2110 |  |  |  |

### Relationship of soil seed bank of the habitats

The relationship of soil seed bank with habitats, piedmont-alluvial (arid-mesic) was analyzed statistically. The analysis confirmed presence of significant differences (p<0.05) of soil seed bank in piedmont and alluvial environment (Table 5) the seed bank increased sequentially from upland inclined gradient (almost non cultivated) to alluvial plains of landscape (cultivated area) (Tables 4 and 5).

**Table 5.** Analysis of variance of 80 different species of seed bank observed from different soil cores depth (0-5, 5-10 and 10-15cm depths) of 2 major different localities of landscape of Dera Ghazi Khan.

| Model 1 | Df | Sum of squares | Mean squares | F-Value | P.Value |
| --- | --- | --- | --- | --- | --- |
| Regression | 1 | 3940.35 | 3940.35 | 9.90 | 0.005*** |
| Residual | 34 | 9434.19 | 398.84 |  |  |
| Total | 35 | 14176.54 |  |  |  |

\*\*Significance at P (0.05)

## DISCUSSION

Seeds which become the part of soil micelles during the seasons, then these seeds were faced many environmental stresses directly or indirectly in the soil for their survival, e.g microbes of soil and harshness of environment is directly effects and destroy the viability nature of seeds present in the soil profile. These evidences were supported by the various international research community like Chesson (1984); Fenner and Thompson (2005); Gioria and Osborne 2009a, (2010). Sometimes the invaders and other related species of wild vegetation community may not form a viable seed bank and their contribution to the soil seed bank for a short duration. Gioria and Osborne (2009a, 2010) were agreed such type of investigation related to invaders species role in terms of seed bank. Moreover, all those species which have bigger and persistent seed bank in the soil, they were poorly represented the above ground vegetation. But in the present investigation the above ground vegetation was the mirror image of the below ground soil seed bank and disagreed with the findings of no relationship of the above ground vegetation and persistent soil seed bank. It was observed that generally the species which produced small and large seeds were having variability to become the part of soil seed bank, thus the seed size and mass play a important role in the persistent seed bank in the soil profile. The trade-off condition of seed mass and size was followed the findings of Thompson *et al*. (1997, 1998). Fenner and Thompson, (2005) were also observed that the seed size and mass of various species, which was a part of soil seed bank reflects with the environmental features of the habitats, such type of observation was also found in the present investigation. The soil seed bank quantification and assessment in the present investigation was link to the habitats and the type of edaphic factors. Edaphic factors more or less were played a role in the preservation and seed viability during the seed bank in the soil profile

In general, the composition and quantification of soil seed bank were totally varied in disturbed and undisturbed habitats, e.g. piedmont soil of hilly mountainous range of intermediate status in the recovery of soil seed bank of the investigated area than the disturbed or regularly cultivated habitats of alluvial plains. These findings were agreed with the previous investigation of Thompson and Grime (1979) and Hopfensberger (2007). A photoblastic and non-photoblastic seeds concept was also difficult in assessing the link of above ground vegetation and below ground vegetation, such type of studies were the findings of Baskin and Baskin (1998). The environmental variables were more or less affected in the regeneration strategies of soil seed bank. It is possible that in an adaptive response to the new disturbance regime introduced by an invasion process, resident species rely more on vegetative propagation and less on seed production for their establishment and spread with effects also on their genetic diversity for species possessing both regenerative strategies (Gioria and Osborne 2010, Fennell et al. 2014).

Quantification of the soil seed bank of individual species or whole seed bank communities presents several challenges, because seed banks are highly variable in space and time according to the findings of Thompson and Grime 1979, Thompson et al. 1997. Spatially, seed banks are generally patchy, with seeds of many species clustering around the mother plant this was the observation of these scientists (Fourie 2008, Gioria and Osborne 2009b), and generally decrease with increases in soil depths (Thompson and Grime 1979, Fenner and Thompson 2005).

Piedmont Soil samples collected from the different sites contained the viable seeds numbers which is lesser than the alluvial plains of current study. The maximum soil seed bank was scrutinized from the alluvial soil of the investigated habitats. Total numbers of 50 different plant species of soil seed banks were recovered from the 60 variable sites of investigated area.

The presence of species in the above ground vegetation does not insure that a seed bank will also be present for that species. Some of the taxa found in the vegetation do not occur in the soil seed bank of the study areas. Generally, the low similarity observations were found between above ground vegetation and presence of seeds in the soil micelles in any agriculture/ arable land ecosystem of the area and such type of studies related with the investigation of Gibson and Looney, (1992), they observed that there was no relationship between soil seed bank and the presence of vegetation on the area.

These results suggested the importance of seed bank in determining the structure and composition of above ground weed vegetation types. So from this study it is concluded that the above ground vegetation is the mirror image of below ground vegetation. In the alluvial plains *Fumaria indica* and *Melilotus albus* were recommended as very abundant species according to analysis of seed bank from different soil core units and germination counts, which were two folds higher than the piedmont soil area of the current study. In the alluvial plain areas almost seed banks recovered of mesic types of species

The significant difference was found in the soil seed bank at different depths levels. The quantitative value of seed bank was successively increased from piedmont soil to alluvial plains of Dera Ghazi Khan Landscape, and depth wise soil seed bank was successively decreased from the range 0-5cm to 10-15cm. The highest numbers of seeds in the soil was found at the depth range of 0-10cm in all three localities of the landscape and this idea is agreed with previous findings of Gulshan et al., (2013) and Li et al., (2017), their findings pointed out the quantity of soil seeds decreased with the increased of depth and disturbance intensity of soil habitats.

From the discussion of observed data, it is concluded that maximum soil seed bank present at the depth of 5-10cm. The 10-15cm depth of soil contained minimum numbers of seeds in their micelles and has 3^rd^ place with reference to soil seed bank. Moreover, it was also concluded that the upland area contains xerophytic types of species soil seed bank. But the alluvial plains the diversity of soil seed bank was contained annual types of vegetation.

In the present investigation above ground vegetation of two different soil habitats is totally variable, because the piedmont zeric habitats dominate with the perennial life span xerophytic species and in other side alluvial habitat abounds with almost annual mesophytic type of species. Abundance and rarity of the species in contrasting environment of landscape of the study area is presented in Table 3. Soil seed bank in respective habitats is also confirmed the similarity of above ground vegetation. The present findings are disagreed the findings of Savadogo et al., (2017). They studied that the soil seed bank was not close relationship with the standing vegetation.

Our current study is the reflection of the findings of Vernooy et al., (2017), according to them the diversity of soil seed bank changes with the changing of climatic and edaphic factors and this type of demonstration was already discussed at different regions by the various researchers across the globe e.g. the Kiziba community of vegetation in terms of soil seed bank in Uganda is a very good example of the strategy of maximizing diversity to respond to climate change.

## CONCLUSION

The most of the xerophytic perennial species found in the piedmont habitat and the analysis of soil seed bank from the micelles of piedmont soil was the confirmation of the presence of above ground vegetation, which was exhibited in the piedmont habitat and piedmont xerophytic species almost absent in the alluvial soil (lowland area) and their soil seed bank also. In contrast to alluvial soil habitat species, which are most having annual life span only present in the alluvial soil not in the piedmont habitat. The current studies will be helpful for the researchers to quantify soil seed bank in the cultivated area for the management of undesirable wild plant species. Our conclusion was disagreed with the findings of Li *et al*., (2017), According to them the results showed that seed density and species all decreased with as human disturbance intensity in the soil seed bank increased. Common species between the soil seed bank and vegetation also decreased as disturbance intensity increased. However, the soil seed bank in a land use type with light or moderate disturbance is still sufficient for vegetation restoration compared with that land use type with severe disturbance. The results would be helpful for vegetation restoration and desertification control in semi-arid regions of northern China, and similar regions all over the world.

